# Neurobeachin controls the asymmetric subcellular distribution of electrical synapse proteins

**DOI:** 10.1101/2022.02.07.479472

**Authors:** E. Anne Martin, Jennifer Carlisle Michel, Jane S. Kissinger, Fabio A. Echeverry, Ya-Ping Lin, John O’Brien, Alberto E. Pereda, Adam C. Miller

## Abstract

The subcellular positioning of synapses and their specialized molecular compositions form the fundamental basis of neural circuits. Like chemical synapses, electrical synapses are constructed from an assortment of adhesion, scaffolding, and regulatory molecules, yet little is known about how these molecules localize at specified subcellular neuronal compartments. Here we investigated the relationship between the autism- and epilepsy-associated gene Neurobeachin, neuronal gap junction channel-forming Connexins, and the scaffold ZO1. Using the zebrafish Mauthner circuit we found Neurobeachin localizes to the electrical synapse independent of ZO1 and Connexins. By contrast, we show Neurobeachin is required postsynaptically for the robust localization of ZO1 and Connexins. We demonstrate Neurobeachin binds ZO1 but not Connexins. Finally, we find Neurobeachin is required to restrict postsynaptic electrical synapse proteins to dendrites. These findings reveal a mechanism for the asymmetric synaptic localization of electrical synapse components providing a basis for the subcellular specialization of neuronal gap junctions.

## Introduction

Neuronal synapses can be separated into two classes of fast transmission, chemical and electrical, each of which is required for neural circuit development and function (Pereda, 2014; Jabeen and Thirumalai, 2018). Chemical synapses are well-studied structures with molecularly diverse asymmetrical junctions, which are known to rely upon highly specialized proteins that orchestrate their localization, assembly, and function (O’Rourke et al., 2012; Südhof, 2012). By contrast, electrical synaptic transmission is achieved via neuronal gap junction channels, which in vertebrates are created by apposed hemichannels of Connexin hexamers that allow for the bidirectional flow of ions and other small molecules. In addition to channels, electrical synapses contain complex biochemistries including cell adhesion, scaffold, and regulatory molecules, yet the mechanisms required to construct neuronal gap junctions remain poorly understood (Martin et al., 2020). Electrical synapses form between all neuronal compartments with each uniquely compartmentalized coupling yielding different effects on activity. For example, dendro-dendritic electrical synapses allow for lateral excitation via the spread of local synaptic potentials between neighboring neurons (Vervaeke et al., 2012). Alternatively, axo-axonic electrical synapses can promote strong, highly synchronized firing with axo-axonic coupling proposed to contribute to fast ripples such as those aberrantly present in patients with epilepsy (Draguhn et al., 1998; Roopun et al., 2010; Curti et al., 2012; Simon et al., 2014; Alcamí and Pereda, 2019). Thus, at a cell biological level, the molecular mechanisms by which electrical synapses are established between specific subcellular compartments are critical for appropriate neural circuit formation and function.

While neurons control the subcellular locations of neuronal gap junctions, the mechanisms by which neurons compartmentalize electrical synapse proteins is not well understood. Recent work in *Caenorhabditis elegans*, which use a different class of molecules for forming gap junctions called Innexins, identified a cAMP-dependent signaling pathway regulating the trafficking of Innexins to distal synaptic sites (Palumbos et al., 2021). The findings support the notion of regulatory mechanisms controlling the trafficking of electrical synapse proteins to distinct subcellular locations, but it is not clear if vertebrate Connexins share related pathways. For chemical synapses, the mechanisms controlling the polarized distribution of synaptic proteins to axons and dendrites are well studied and require the coordination of motor proteins, the cytoskeleton, and numerous adaptors and regulators to ensure the fidelity of compartmentalized delivery (Ou et al., 2010; Easley-Neal et al., 2013; Maeder et al., 2014; Mignogna and D’Adamo, 2017). Whether there are distinct, or shared, mechanisms to guide the polarized transport of electrical synapse proteins remains unknown.

The formation of electrical and chemical synapses are generally thought to be cell biologically and biochemically distinct processes. Yet in recent years molecules have emerged that suggest links between the development and function of these structures (Pereda, 2014; Miller et al., 2015; Jabeen and Thirumalai, 2018; Martin et al., 2020). One of these proteins, Neurobeachin, was found to be required for both electrical and chemical synapse development (Miller et al., 2015). Neurobeachin is a large (~330 kDa) protein that is highly conserved in vertebrates and contains various protein-binding domains including a BEACH domain, AKAP domain, and multiple WD40 domains (Wang et al., 2000). More than 20 *de novo* variants of Neurobeachin have been identified in patients with autism, intellectual disability, and epilepsy (De Rubeis et al., 2014; Iossifov et al., 2014; Bowling et al., 2017; Mulhern et al., 2018). Previous work suggests Neurobeachin regulates the localization of chemical synapse proteins including glutamate receptors and the scaffolds SAP102 and PSD95 (Wang et al., 2000; Medrihan et al., 2009; Niesmann et al., 2011; Nair et al., 2013; Farzana et al., 2016). Yet its mechanistic role in electrical synapse development is unknown.

Here we explore the contributions of Neurobeachin to electrical synapse formation and examine its biochemical and cell biological functions. We used the stereotyped and identifiable synaptic contacts of the Mauthner cell of larval zebrafish as they are accessible to genetic, biochemical, and cell biological analysis. We show that Neurobeachin is localized to the electrical synapse and functions postsynaptically to promote the robust localization of the electrical synapse scaffold Zonula Occludens 1 (ZO1) and neuronal Connexins. We find that Neurobeachin interacts with ZO1 but not the Connexin proteins, supporting a hierarchical model of electrical synapse formation. Further, we find that Neurobeachin functions to restrict postsynaptic electrical synapse proteins to dendritic synapses, such that in its absence postsynaptic proteins are inappropriately localized and stabilized at presynaptic locations in the axon. These results reveal an expanded understanding of electrical synapse molecular complexity and the hierarchical interactions required to build neuronal gap junctions. Further, these findings provide novel insight into the mechanisms by which neurons compartmentalize the localization of electrical synapse proteins and provide a cell biological mechanism for the subcellular specificity of electrical synapse formation and function.

## Results

### Neurobeachin is required for the localization of pre- and postsynaptic electrical synapse proteins

To investigate how Neurobeachin regulates electrical synapse development we used the zebrafish Mauthner cell circuit, which uses a combination of electrical and chemical synapses to control a fast escape response (Kimmel et al., 1981; Liu and Fetcho, 1999; Sillar, 2009). This circuit is composed of two Mauthner cells per fish that receive sensory input from several modalities, including the auditory system, and send output onto motorneurons and interneurons in the spinal cord, including Commissural Local (CoLo) interneurons (Figure 1A,B). Electrical synapses are prominent throughout the circuit, including at mixed electrical/glutamatergic inputs from auditory afferents onto the lateral Mauthner dendrite, called club ending (CE) synapses (Yao et al., 2014). Additionally, the Mauthner cell uses electrical synapses between its axon and CoLos, called M/CoLo synapses (Figure 1B). These M/CoLo synapses repeat down the length of the spinal cord, with CoLo being present as a single cell in each of the ~30 hemisegments. Mauthner and CoLo cell morphology can be observed using the transgenic line *Et(Tol-056:GFP)(Satou* et al., 2009). The CE and M/CoLo electrical synapses are molecularly asymmetric composed of presynaptic Connexin 35.5 hemichannels (Cx35.5 – encoded by the *gjd2a* gene) that pair with postsynaptic Cx34.1 hemichannels (encoded by the *gjd1a* gene) (Miller et al., 2017). In addition, the electrical synapse scaffold ZO1b (encoded by the *tjp1b* gene) localizes exclusively postsynaptically where it binds Cx34.1 (Figure 1B) (Marsh et al., 2017; Lasseigne et al., 2021). At the CE synapses, the Mauthner cell is the postsynaptic partner and uses ZO1b and Cx34.1 to build electrical synapses, while at the M/CoLo synapses, the Mauthner cell is presynaptic and uses Cx35.5 (Figure 1B). The asymmetry of Mauthner cell electrical synapses highlights the cell biological requirement of the neuron to deliver unique electrical synaptic proteins to distinct neuronal compartments.

**Figure 1:**
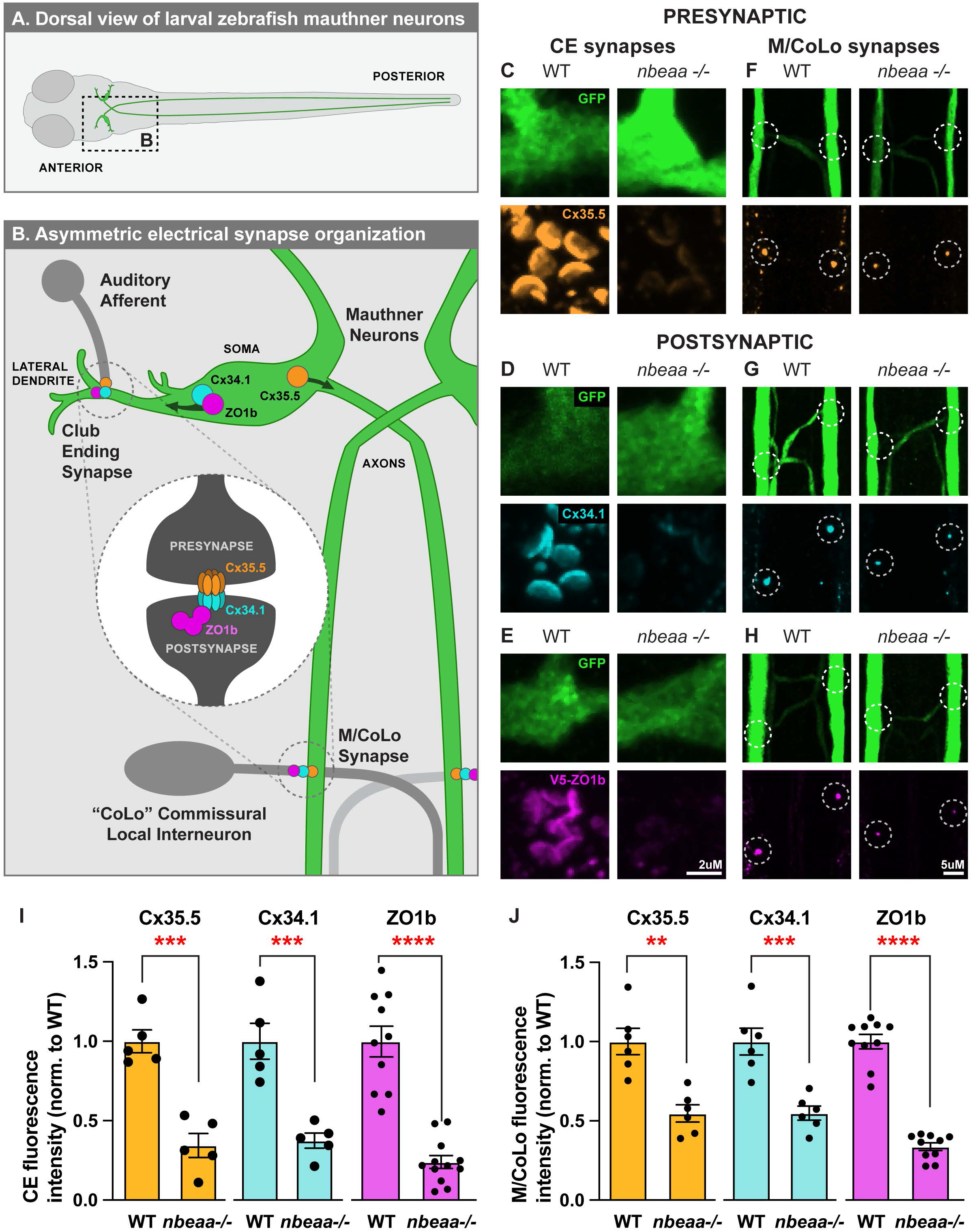
Neurobeachin is required for the localization of pre- and postsynaptic electrical synapse proteins. **(A)** Dorsal view of the larval zebrafish illustrating location of the two Mauthner neurons (green), anterior left. Boxed region indicated the region shown in B. **(B)** Simplified illustration of the Mauthner circuit, anterior up, which identifies the synapses examined and the known protein localization of electrical synapse components Cx35.5 (orange), Cx34.1 (cyan), and ZO1b (magenta). The center dashed-circle diagram depicts the standard synapse organization for each Mauthner synapse which includes gap junction hemichannels composed of Cx35.5 in the presynapse, and Cx34.1 in the postsynapse where it binds scaffolding protein ZO1b. The top left dashed circle along the Mauthner lateral dendrite indicates a Club Ending (CE), which connects an auditory afferent to Mauthner. The bottom right dashed circle along the Mauthner axon indicates an M/CoLo synapse, which connects Mauthner to Commissural Local (CoLo) interneurons. These axonal synapses are continued pairwise, in each segment of the fish moving down the spinal cord. Arrows in Mauthner soma indicate the direction of subcellularly localized of postsynaptic proteins Cx34.1 and ZO1b to the CE synapses in the dendrite, and presynaptic Cx35.5 to the M/CoLo synapses in the axon. **(C-H)** Confocal images of Mauthner synapses in 5 dpf *Et(Tol-056:GFP)*, transgenic *wildtype (WT)* or *nbeaa*-/- zebrafish. Panels show staining for GFP labeling the Mauthner cell (green), Cx35.5 (orange), Cx34.1 (cyan), V5-ZO1b (magenta) at: **(C-E)** CE synapses, and **(F-H)** M/CoLo synapses. Dashed circles identify locations of Mauthner and CoLo contact. **(I-J)** Quantification of indicated electrical synapse protein fluorescence intensities, normalized to WT, at (I) CE synapses and (J) M/CoLo synapses in WT and *nbeaa*^−/−^ mutant animals. Mean ± SEM are shown, All comparisons were made between genotyped WT and mutant siblings. For I, n = 5, 5, 5, 5, 10, and 12 fish for each bar shown, left to right. Two Mauthner neurons were analyzed per fish and averaged to represent a single value for each fish. For graph in J, n = 6, 6, 6, 6, 10, and 10 fish, for each bar shown, left to right. 20-24 M/Colo synapses were analyzed and averaged for each fish. ** indicates p < 0.01, *** indicates p < 0.001, **** indicates p < 0.0001 by unpaired t-test. Scale bars are as indicated.

We first examined the role of Neurobeachin (encoded by the *nbeaa* gene) in localizing the presynaptic Cx35.5 and the postsynaptic Cx34.1 and ZO1b to Mauthner cell electrical synapses (Figure 1C-J). We previously found that Neurobeachin is required for Connexin localization to Mauthner cell electrical synapses (Miller et al., 2015), however, the molecular identity of the Connexins, the ZO1b scaffold, and the asymmetry of these proteins at the synapse were unknown at the time. We examined 5 days post fertilization (dpf) wildtype and *nbeaa^fh364/fh364^* mutants (hereafter called *nbeaa*-/-; Miller et al., 2015), for Cx35.5 and Cx34.1 localization at CE (Figure 1C,D,I) and M/CoLo synapses (Figure 1F,G,J). In *nbeaa*-/- mutants, we observed similar decreases of both pre- and postsynaptic Connexin proteins at synapses. Using a transgenic line that marks endogenous ZO1b with a V5 epitope (*tjp1b^b1406^*, hereafter called *Pt(V5-ZO1b)*, Lasseigne et al., 2021), we examined 5 dpf wildtype and *nbeaa*-/- mutant fish for ZO1b at CE (Figure 1E,I) and M/CoLo synapses (Figure 1H,J). We observe a similar loss of ZO1b at Mauthner dendritic and axonal electrical synapses in *nbeaa*-/- mutant fish. The diminished synaptic localization is likely not due to a defect in neurite outgrowth as wildtype and *nbeaa*-/- Mauthner cell dendrites and axons form normally and do not show altered targeting or gross morphological defects (Supplemental Figure 1A). Additionally, western blots confirm similar Cx35.5, Cx34.1, and ZO1b protein levels are present in both wildtype and *nbeaa*−/− mutant fish (Supplemental Figure 1B), supporting the notion that mutants have defects in robust localization of electrical synapse proteins. We conclude that Neurobeachin is required for the synaptic localization of pre- and postsynaptic electrical synapse proteins.

### Neurobeachin localizes to electrical synapses independent of ZO1 and Connexins

In electron microscopic analyses of rat cerebellar neurons, Neurobeachin localizes to the trans side of the Golgi, on tubulovesicular vesicles in dendrites, and at the postsynapse of glutamatergic chemical synapses (Wang et al., 2000). Thus, we next examined Neurobeachin localization at electrical synapses of the Mauthner cell circuit. At 5 dpf, we observe Neurobeachin localization in the Mauthner cell soma, but little colocalization with electrical synapse proteins in this compartment (Figure 2A). To examine Neurobeachin’s somatic distribution, we stained for the Golgi apparatus (GM130) and observed the Mauthner cell of wildtype and *nbeaa*−/− mutants. Golgi staining within the Mauthner soma was found throughout the compartment but asymmetrically biased towards the axon; this distribution was not altered in *nbeaa*−/− mutants (Supplemental Figure 2A). In addition, we noticed a high degree of close-proximity localization of Neurobeachin with GM130, similar to what has been observed in rodents (Wang et al., 2000; Nair et al., 2013).

**Figure 2:**
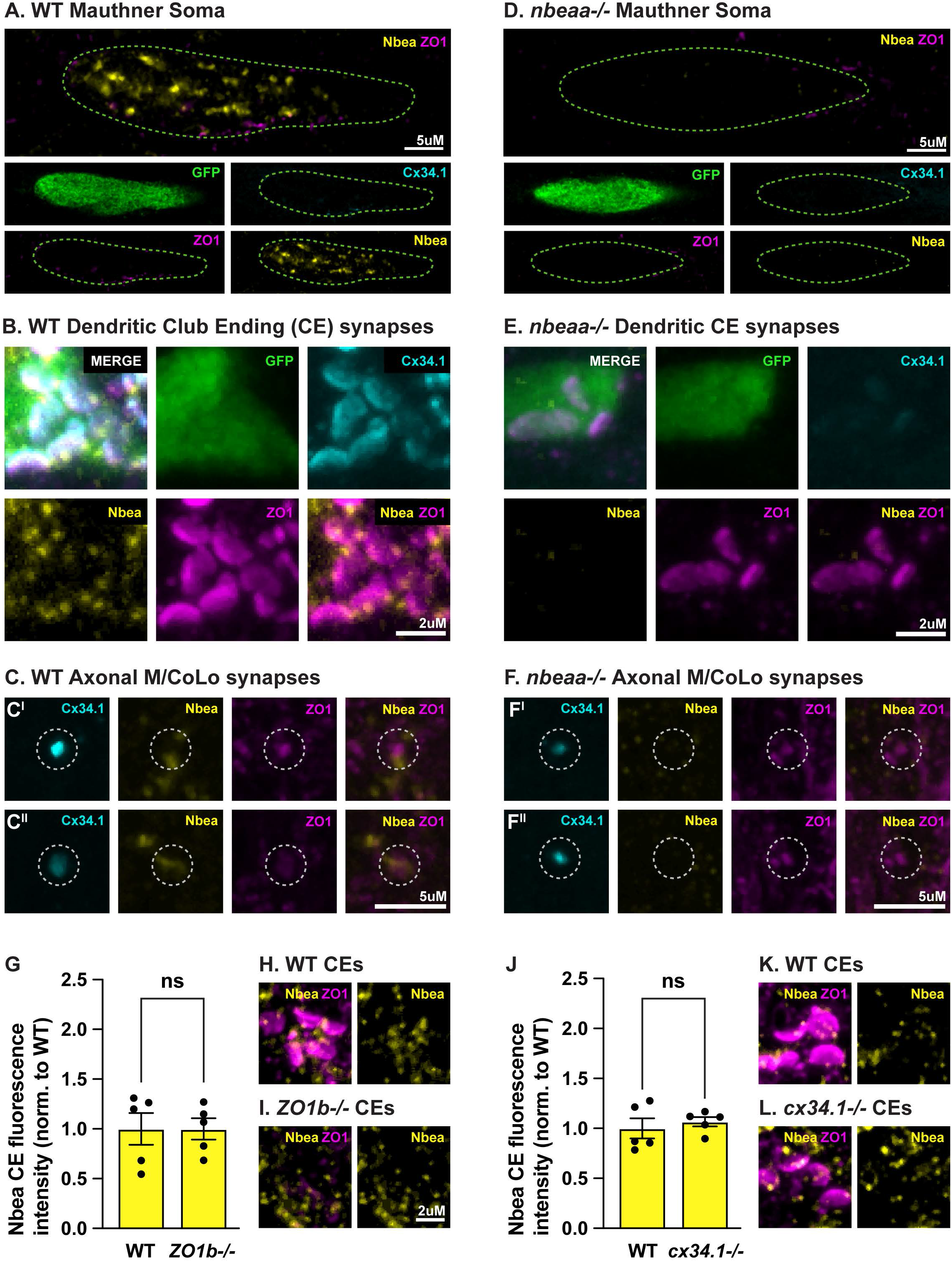
Neurobeachin localizes to the electrical synapse independent of ZO1b and Connexin proteins. **(A-F)** Confocal images of Mauthner subcellular regions in 5-days-post-fertilization (dpf) *Et(Tol-056:GFP)* transgenic *wildtype* (WT, A-C) and *nbeaa*−/− (D-F) zebrafish, as noted. Panels show staining for GFP labeling the Mauthner cell (green), Cx34.1 (cyan), ZO1 (magenta), and Neurobeachin (yellow) at: **(A,D)** Mauthner cell soma, **(B,E)** dendritic CE synapses, and **(C,F)** axonal M/CoLo synapses. Images are maximum intensity projections. Green dashed outline in A represents the cell soma. White dashed outlines in C,F represent the M/CoLo electrical synapse site. C^I^, C^II^ and F^I^, F^II^ represent two distinct M/CoLo synapses for each genotype. **(G-L)** Neurobeachin localization at Mauthner CE synapses in 5 dpf *Et(Tol-056:GFP)* transgenic zebrafish from *wildtype* (WT, H,K), *tjp1b/ZO1b*^−/−^ mutant animals (I), and gjd1a/Cx34.1^−/−^ mutant animals (L). **(G)** Quantification of Neurobeachin fluorescence intensities at *wildtype* (WT) and *tjp1b/ZO1b*^−/−^ (ZO1b−/−) CE normalized to WT. **(H,I)** Representative images of wildtype (H) and *tjp1b/ZO1b*^−/−^ (ZO1b−/−) (I) CE synapses with markers as indicated. **(J)** Quantification of Neurobeachin fluorescence intensities at WT and *gjd1a/Cx34.1*^−/−^ (Cx34.1−/−) CE synapses normalized to WT. **(K,L)** Representative images of wildtype (K) and *gjd1a/Cx34.1*^−/−^ (Cx34.1−/−) (L) CE synapses with markers as indicated. Circles in the bar graphs represent the normalized value of each individual animal (CE synapses from 2 dendrites per animal averaged to produce single value for each animal). For G-L, comparisons were made between genotyped WT and mutant sibs. *wildtype* (H) n = 5, *wildtype* (K) n = 5, *tjp1b/ZO1b*^−/−^ (I) n = 5, *gjd1a/Cx34.1*^−/−^ (L) n = 5. No significant difference found by unpaired t-test. Error bars are ± SEM. Scale bars are as indicated.

At the CE synapses on Mauthner’s lateral dendrite, we see Neurobeachin localization at distinct foci adjacent to the electrical synapse proteins ZO1b and Cx34.1, as well as a diffuse staining colocalized across the extent of the observed gap junction staining (Figure 2B). We also observe Neurobeachin staining at M/CoLo synapses, with staining that always appears to be adjacent to ZO1 and Connexins, and we never observe staining within the prominent Mauthner axon between each synapse (Figure 2C). We note that at each Mauthner associated synapse (CE or M/CoLo) the pre- or postsynaptic localization of Neurobeachin cannot be resolved due to the limits of light microscopy — we address where the protein functions below. Neurobeachin staining is eliminated in *nbeaa*−/− mutants supporting the specificity of the antibody (Figure 2D-F, Supplementary Figure 2B). We conclude that Neurobeachin can localize to electrical synapses, yet does so in a manner distinct from the scaffold and neural Connexins.

Next, we examined whether Neurobeachin localization to the electrical synapse was dependent upon the function of other electrical synapse proteins. The intracellular scaffold ZO1b is required for Connexin localization, but ZO1b largely localizes to electrical synapses independently of the Connexins (Lasseigne et al., 2021). We therefore examined Neurobeachin localization in ZO1b and Connexin mutants. We examined *ZO1b/tjp1b^b1370/b1370^* (Marsh et al., 2017) and *Cx34.1/gjd1a ^fh436/fh436^* (Miller et al., 2017) mutants and found that Neurobeachin localizes to electrical synapses in a manner similar to their wildtype siblings (Figure 2G-L). Overall, we conclude that Neurobeachin localizes to the electrical synapse independent of ZO1b and Connexins.

### Neurobeachin binds the electrical synapse scaffold ZO1b but not the Connexins

Neurobeachin is thought to orchestrate the delivery of receptors to the glutamatergic chemical synapse through binding with chemical synapse scaffolds such as membrane-associated guanylate kinase (MAGUK) family members PSD95 and SAP102 (Lauks et al., 2012; Farzana et al., 2016). Given that the electrical synapse scaffold ZO1b is also a member of the MAGUK family, we next asked if there was a Neurobeachin/ZO1b interaction. We attempted to clone full-length *nbeaa* sequences from fish, but the size prevented the recovery of the entire coding sequence. We were able to generate N- and C-terminal fragments of Neurobeachin, each tagged with mVenus (Fig. 3A), and used heterologous expression in HEK293T cells to test for interactions with previously cloned zebrafish ZO1b and neural Connexins (Lasseigne et al., 2021). We co-transfected mVenus-tagged C-terminal Neurobeachin with ZO1b and performed coimmunoprecipitation assays. We find that the C-terminal fragment of Neurobeachin pulls down with ZO1b (Figure 3B). By contrast, we find no evidence for interaction between the N-terminal fragment of Neurobeachin and ZO1b (Supplementary Figure 3A). We next tested whether Neurobeachin could bind the pre- or postsynaptic Connexins. We co-transfected each Connexin with N-terminal and C-terminal Neurobeachin and find no evidence of pull down with Neurobeachin in immunoprecipitates (Figure 3C,D). By contrast, we find robust interaction between ZO1b and both Connexins, as previously shown (Lasseigne et al., 2021). We conclude that Neurobeachin can bind the postsynaptic electrical synapse scaffold ZO1b.

**Figure 3:**
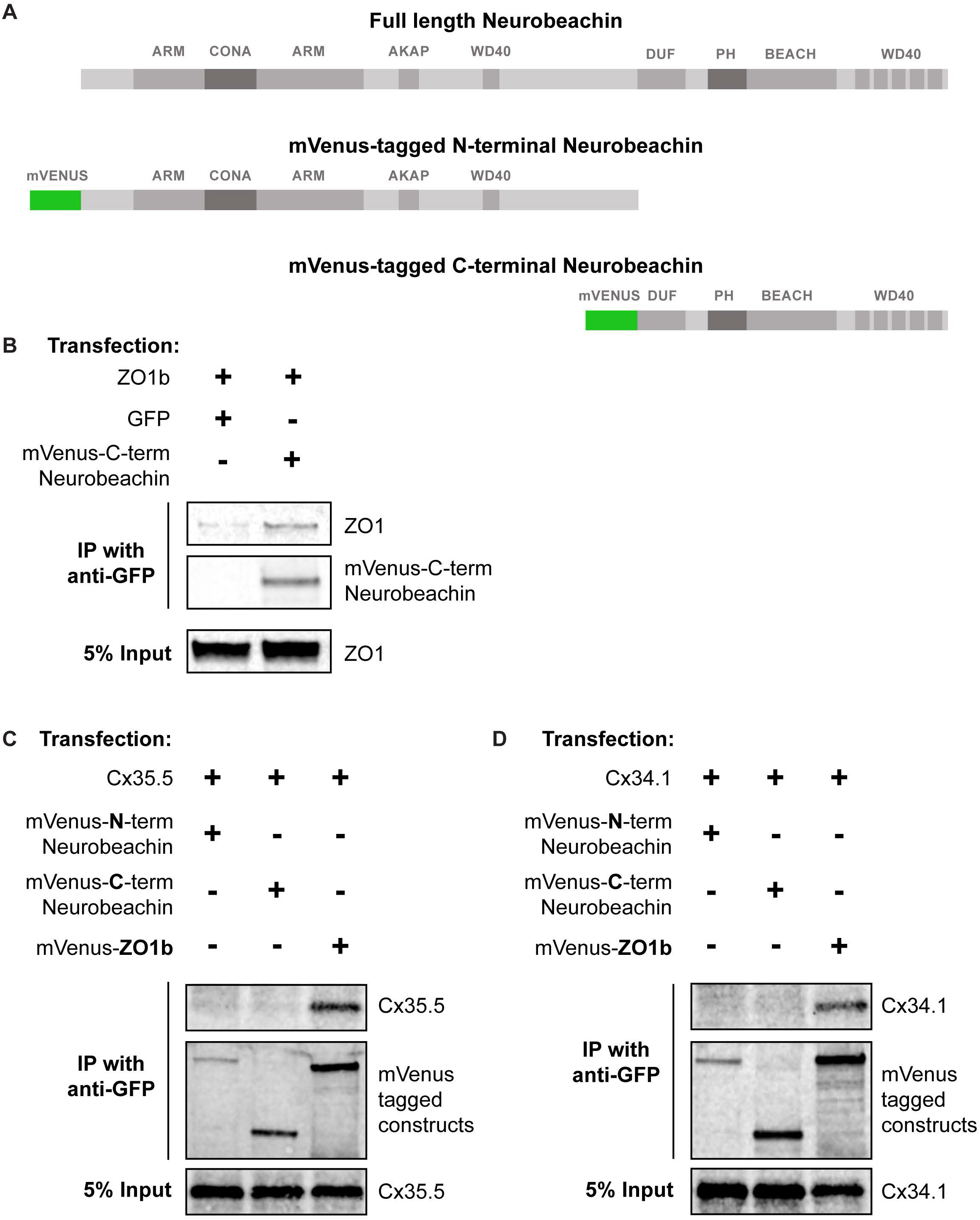
Neurobeachin binds the postsynaptic electrical synapse scaffold ZO1b, but not the neural Connexins. **(A)** Schematic of Neurobeachin protein indicating domains included in mVenus-tagged N- and C-terminal fragments. **(B)** HEK293T/17 cells were transfected with plasmids to express ZO1b and either GFP (lane 1), or mVenus-tagged C-terminal-Neurobeachin (lane 2). Lysates were immunoprecipitated with anti-GFP antibody and analyzed by immunoblot for the presence of ZO1b using ZO1 antibody (upper), or mVenus-tagged C-terminal Neurobeachin using GFP antibody (middle). Total extracts (bottom, 5% input) were blotted for ZO1b to demonstrate equivalent expression and uniform antibody recognition of expressed proteins. **(C)** HEK293T/17 cells were transfected with plasmids to express Cx35.5 and either mVenus-tagged N-terminal Neurobeachin (lane 1), mVenus-tagged C-terminal-Neurobeachin (lane 2), or mVenus-tagged ZO1b (lane 3). Lysates were immunoprecipitated with anti-GFP antibody and analyzed by immunoblot for the presence of Cx35.5 using a specific Cx35.5 antibody (top), mVenus-tagged N-terminal Neurobeachin using GFP antibody (middle left), mVenus-tagged C-terminal Neurobeachin using GFP antibody (middle middle), or mVenus-tagged ZO1b using GFP antibody (middle right). Total extracts (bottom, 5% input) were blotted for Cx35.5 to demonstrate expression. **(D)** HEK293T/17 cells were transfected with plasmids to express Cx34.1 and either mVenus-tagged N-terminal Neurobeachin (lane 1), mVenus-tagged C-terminal-Neurobeachin (lane 2), or mVenus-tagged ZO1b (lane 3). Lysates were immunoprecipitated with anti-GFP antibody and analyzed by immunoblot for the presence of Cx34.1 using a specific Cx34.1 antibody (top), mVenus-tagged N-terminal Neurobeachin using GFP antibody (middle left), mVenus-tagged C-terminal Neurobeachin using GFP antibody (middle middle), or mVenus-tagged ZO1b (middle right). Total extracts (bottom, 5% input) were blotted for Cx34.1 to demonstrate expression.

### Neurobeachin is required postsynaptically for pre- and postsynaptic electrical synapse protein localization

Observing that Neurobeachin binds ZO1b, which localizes and functions postsynaptically (Lasseigne et al., 2021), we next wanted to examine whether Neurobeachin also functioned postsynaptically *in vivo*. To achieve this we generated chimeric animals via blastula transplantation (Kemp et al., 2009) (Figure 4A) and we first examined the localization of ZO1b within the Mauthner cell. Donor embryos were *Et(Tol-056:GFP)*; *Pt(V5-ZO1b)* double transgenics, which together label Mauthner cells with GFP and mark the endogenous ZO1b protein with a V5 epitope tag. All host embryos were non-transgenic wildtypes – thus the observed V5-ZO1b staining is associated exclusively with the transplanted GFP+ Mauthner donor cells. The double transgenic donor embryos were either *nbeaa+/+* (WT) or *nbeaa*−/− mutant, allowing us to examine the effects of the loss of Neurobeachin from the Mauthner cell. In *nbeaa*−/− mutant Mauthner cells, localization of V5-ZO1b at the dendritic CE synapses was reduced as compared to wildtype transplants (Figure 4B), revealing Neurobeachin is required postsynaptically for ZO1 localization to the synapse. In analogous transplants, we next examined the effect of removing Neurobeachin from Mauthner on Connexin localization. For the transplant donor cells, we used the *Tg(Cx34.1-GFP^AE37^)* transgenic line that labels Cx34.1 with GFP. At the dendritic CEs of *nbeaa*−/− mutant Mauthner neurons, we observe that the localization of postsynaptic Cx34.1-GFP and presynaptic Cx35.5 (labeled by antibody) are both reduced as compared to wildtype transplants (Figure 4C). These data show that the removal of Neurobeachin exclusively from the postsynaptic neuron affects the localization of the dendritically localized ZO1b and Cx34.1. In addition, the removal of Neurobeachin from Mauthner affects presynaptic Cx35.5 localization in the neighboring auditory afferent neuron. Taken together, the data support an expanded, asymmetric, electrical synapse hierarchy, with Neurobeachin independently localizing to neuronal gap junctions where it binds ZO1b and is required autonomously for postsynaptic ZO1/Connexin localization, and non-autonomously for presynaptic Connexin localization.

**Figure 4:**
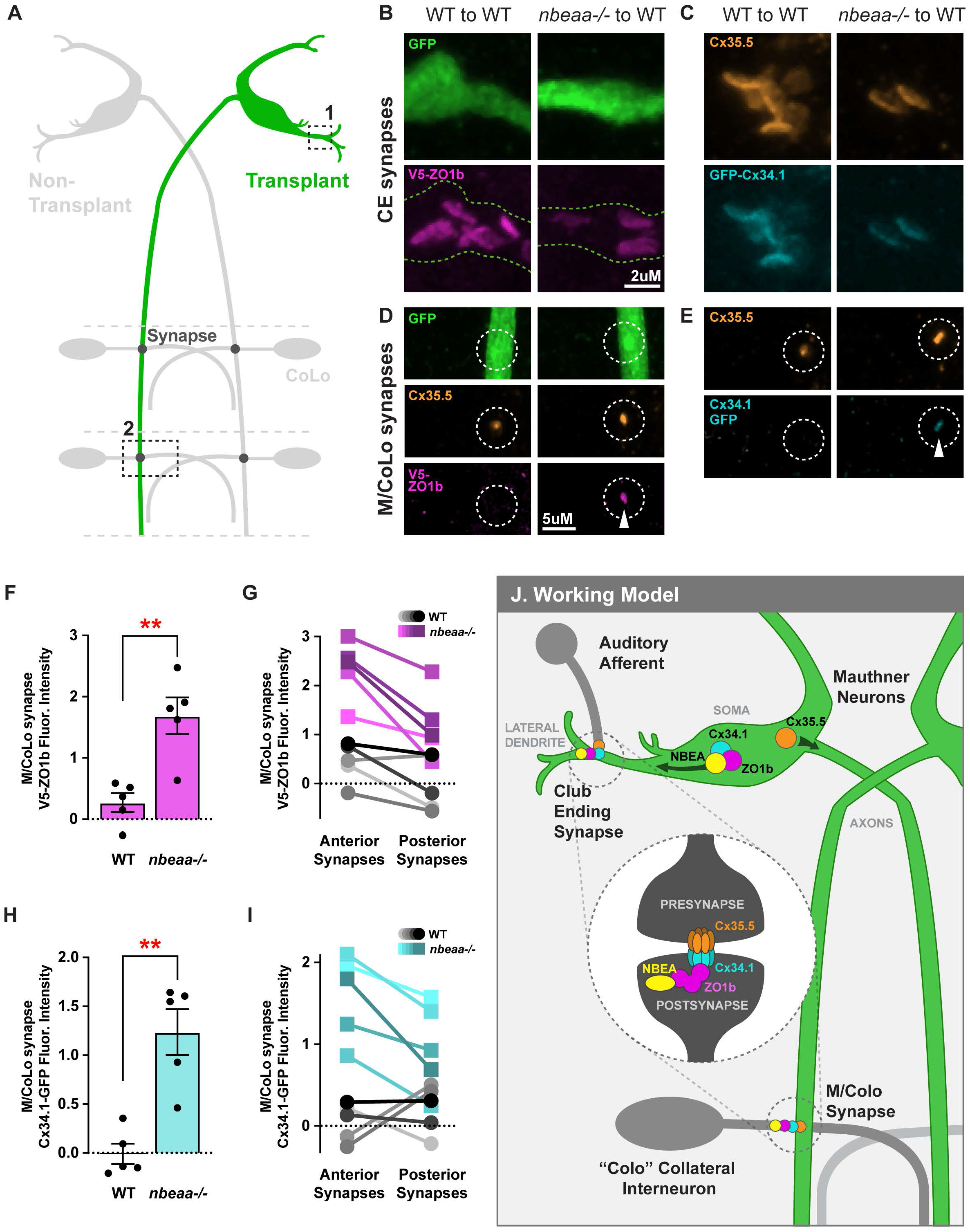
Neurobeachin is required postsynaptically for the proper dendritic compartmentalization of ZO1b and Cx34.1. **(A)** Diagram illustrating 5 dpf Mauthner circuit after transplantation of a single Mauthner neuron (green). Boxed outline 1: Location of CE synapses shown by representative confocal images in B and C. Boxed outline 2: Location of M/CoLo synapses shown by representative confocal images in D and E. **(B,D)** Confocal images of chimaeric animal in which Mauthner is derived from *Et(Tol-056:GFP); Pt(V5-ZO1b)* double transgenic animals, while the rest of the cells are wildtype (WT). Mauthner CE synapses (B) and M/CoLo synapses (D) identified by GFP-positive Mauthner cell. Left images show transplanted WT Mauthner cell (GFP+) in a WT host with associated postsynaptic V5 staining for V5-ZO1b produced from transplanted cell. Right images show a transplanted *nbeaa*−/− Mauthner cell in an WT host with reduced V5-ZO1b at CE synapses (B) and with associated accumulation of staining at M/CoLo synapses (D). Outline of Mauthner cell shown by green dashed line in (B,C). M/CoLo sites of contact shown by white dashed circles in (D,E). At M/CoLo synapse, Cx35.5 was also stained. White arrowhead shows aberrant presynaptic ZO1b. **(C,E)** Confocal images of chimaeric animal in which Mauthner is derived from Cx34.1-oxGFP transgenic animals, while the rest of the cells are wildtype (WT). Mauthner CE synapses (C) and M/CoLo synapses (E) are identified by stereotypical Cx35.5 staining. Left images show transplanted WT Mauthner cell (Cx34.1-GFP+) in an WT host with staining for Cx34.1-GFP produced from transplanted cell. Right images show a transplanted *nbeaa*−/− Mauthner neuron in an WT host with reduced Cx34.1-GFP at CE synapses (C) and associated accumulation of staining at M/CoLo synapses (E). White arrowhead shows aberrant presynaptic Cx34.1. **(F-I)** Quantification of labeling at all M/CoLo synapses in wildtype Mauthner neuron transplants (WT) and in *nbeaa*−/− Mauthner cell transplants for V5-ZO1b (F,G) or Cx34.1-GFP (H,I). Relative fluorescence intensity is quantified and graphed based on fold change normalized to expression at non-transplanted synapses in the same animal. (G,I) Anterior segments (first eight segments imaged) and posterior segments (next eight segments imaged) were quantified and averaged for fluorescence as noted above. **(J)** Diagram showing proposed Neurobeachin (yellow) roles in regulating Mauthner neuron electrical synapse proteins. Mean ± SEM are shown, All comparisons were made between genotyped WT and mutant siblings. For (F,G) *wt*, n = 5, *nbeaa*−/−, n = 5. For (H,I) *wt*, n = 5, *nbeaa*−/−, n = 5. ** indicates p < 0.01 by unpaired t-test. Scale bars are as indicated.

### Neurobeachin is required to restrict postsynaptic electrical synapse proteins to the dendritic compartment

We next took advantage of the fact that the Mauthner cell is both a postsynaptic partner at the CEs as well as the presynaptic partner at the M/CoLo electrical synapses. This configuration, in conjunction with the chimeric transplant experiments, allows us to visualize the compartmentalized localization of components of the electrical synapse. In wildtype transplants, the V5-tagged ZO1b, which is expressed exclusively from the transplanted Mauthner cell in these embryos, is exclusively localized at CE synapses and is not present at M/CoLo synapses (Figure 4B,D, left panels). Thus, within the Mauthner cell, ZO1b is compartmentalized exclusively postsynaptically to the dendritic synapses and excluded from localizing presynaptically within the axon. Strikingly, in *nbeaa*−/− transplanted Mauthner cells, we observed localization of V5-ZO1b presynaptically at M/CoLo synapses (Figure 4D, right panels, quantified in F). The Mauthner cell axon extends the length of the spinal cord and it makes *en passant* electrical synapses with CoLo neurons found in each hemi-segment. The mislocalized V5-ZO1b appears strongly at M/CoLo synapses found in the anterior half of the spinal cord, and diminishes at synapses found posteriorly (Figure 4G). We conclude that Neurobeachin is required to restrict ZO1b localization to the dendritic compartment and prevent it from localizing at axonal electrical synapses.

We next questioned whether the loss of Neurobeachin from the Mauthner cell would affect Connexins. In control transplants, Cx34.1-GFP is exclusively found compartmentalized to the CE synapses and is excluded from M/CoLo synapses (Figure 4C,E, left panels); this is the same pattern as observed with ZO1b (Fig. 4B,D). By contrast, we observe that *nbeaa*−/− Mauthner neurons mislocalize Cx34.1 presynaptically at axonal M/CoLo synapses (Figure 4E, right panels, quantified in H). In addition, analogous to our V5-ZO1b transplant experiments, this mislocalization of Cx34.1 appears strongly at anterior M/CoLo synapses and diminishes posteriorly (Figure 4I). We note that the removal of *nbeaa−/− from* Mauthner cells does not affect Cx35.5 staining at M/CoLo synapses as compared to wildype (Fig. 4E) – this suggests that Neurobeachin does not function presynaptically to localize Connexins. To test this directly, we examined whether presynaptic Cx35.5 would mislocalize to dendritic postsynaptic CE synapses in *nbeaa*−/− mutants (Supplementary Figure 4). We used CRISPR to V5-tag endogenous Cx35.5 protein and examined injected mosaic F0 animals for the localization of Cx35.5-V5 in wildtype and *nbeaa*−/− animals. In wildtype, we identified injected animals in which all M/CoLo synapses have V5 staining, as expected for the presynaptic usage of Cx35.5 in the Mauthner axon; in such cases, no Cx35.5-V5 staining was observed at CE synapses (Supplementary Figure 4A,B). In *nbeaa*−/− mutants, we found animals with the same axonal Cx35.5-V5 present at all M/CoLo synapses (Supplementary Figure 4D), however the amount localized was reduced as expected in *nbeaa*−/− mutants (compare to Figure 1C,F). Critically, in such animals, no Cx35.5-V5 staining is observed at CE synapses (Supplementary Figure 4C). We conclude that Neurobeachin is not required for axonal Cx35.5 compartmentalization, but is required to restrict Cx34.1 to the dendrite.

Taken together, we find that in the absence of Neurobeachin, both the postsynaptic ZO1b scaffold and postsynaptic Cx34.1 are reduced at the dendritic CEs, and instead mislocalize to the axonal M/CoLo presynapses. By contrast, the presynaptic Cx35.5 is unaffected by the loss of Neurobeachin. Thus, we conclude that Neurobeachin functions specifically to constrain dendritically localized synaptic components to their appropriate compartment, thereby supporting the subcellular specificity of electrical synapse formation (Figure 4J).

## Discussion

The experiments described here investigating Neurobeachin provide new biochemical and cell biological insight into the development of electrical synapse molecular organization. First, the findings expand the understanding of the hierarchical construction of the molecular components of the electrical synapse. We find that Neurobeachin localizes to the electrical synapse independent of ZO1b or Connexins, yet Neurobeachin is necessary for these proteins to robustly localize to the developing synapse. Furthermore, we have identified a biochemical connection between Neurobeachin and ZO1b, and previous work found that ZO1b in turn binds Cx34.1 (Lasseigne et al., 2021). Thus, our emerging model is that Neurobeachin sits at the top of an electrical synapse hierarchy, directing ZO1b to the synapse, which in turn brings in the Connexin to build the neuronal gap junction. We further show that Neurobeachin functions postsynaptically in building the electrical synapse, and so why are presynaptic Cx35.5 proteins affected? We have observed that the removal of postsynaptic Cx34.1 from the synapse results in a decrease in presynaptic Cx35.5 localization (Miller et al., 2017). Thus, we hypothesize that the decreased presynaptic Cx35.5 observed in *nbeaa*−/− animals is most likely due to the diminished postsynaptic Cx34.1 localization. These results add further support to the notion that electrical synapses, like chemical synapses, are molecularly complex synaptic compartments, with asymmetric biochemical compositions that have both autonomous and non-autonomous effects on building the electrical synapse.

The results presented here establish that Neurobeachin is required for the compartmentalized synaptic localization of postsynaptic electrical synapse proteins. Given that Neurobeachin localizes on vesicular structures near the Golgi, in the dendrites, and also at electrical synapses, it is likely that it directs synaptic components to the dendrite. Within the Mauthner cell, we are unable to observe Connexin and ZO1b proteins while they are being trafficked to their synaptic sites. Instead, at this time, we can only view these proteins once they have accumulated at synaptic contacts. Therefore, we cannot exclude the possibility that in wildtype neurons, electrical synapse proteins are trafficked into each compartment and are only stabilized at appropriate synaptic contacts by Neurobeachin. Future technical improvements in detection and imaging will be required to assess the trafficking pathways taken by these, and other, electrical synapse proteins. We note that in *nbeaa*−/− mutants, we observe normal levels electrical synapse proteins abundance (by western blot), as compared to wildtype, suggesting the proteins are present but not appropriately localized to the synapse. Despite this, our results clearly demonstrate that Neurobeachin is required to restrict postsynaptic ZO1b and Cx34.1 to their normal dendritic synaptic locations. Taken together, we propose a model in which Neurobeachin is necessary for the trafficking and/or stabilization of postsynaptic proteins to the electrical synapse (Figure 4J).

Why would a neuron compartmentalize electrical synapse components? The localization of electrical synapses at distinct subcellular compartments has critical impacts on neuronal function, such as lateral excitation via dendro-dendritic electrical synapses (Vervaeke et al., 2012) or strong synchronization via axo-axonic electrical synapses (Draguhn et al., 1998; Roopun et al., 2010; Curti et al., 2012; Simon et al., 2014). These configurations require the compartmentalized delivery or capture of electrical synapse components to unique sites. In the Mauthner cell, such compartmentalization creates molecularly asymmetric electrical synapses, with unique pre- and postsynaptic biochemistries (Marsh et al., 2017; Miller et al., 2017). Molecular asymmetry is an important feature of electrical synapses which can determine functional properties, for instance to allow for electrical rectification (Rash et al., 2013). In *nbeaa*−/− animals, postsynaptic proteins are inappropriately localized presynaptically, along with the normal presynaptic proteins, at the axonal presynapse. That means that these synapses have become more symmetric, with ZO1 and Cx34.1 on both sides of the synapse, which may then alter the functional properties of the circuit. Thus, Neurobeachin represents a cell biological mechanism by which neurons can direct electrical synapse components to appropriate locations, building unique biochemistries at subcellular localizations to create specific circuit functions.

How might Neurobeachin direct electrical synapse proteins to specific compartments? Neurobeachin contains an A-Kinase Anchoring Protein (AKAP) domain which could promote the formation of a synaptic PKA signaling complex to phosphorylate electrical synapse proteins upon cAMP stimulation (Wang et al., 2000; Autenrieth et al., 2016). Phosphorylation can alter gap junction processes such as trafficking, assembly, degradation, and channel function (Lampe and Lau, 2000). Given the findings in C. elegans that cAMP signaling is necessary for the trafficking of Innexin proteins to specific subcellular regions (Palumbos et al., 2021), it is enticing to speculate that Neurobeachin may be the cell biological link between cAMP signaling and the compartmentalized trafficking of electrical synapse components. Overall, these overlapping pieces of evidence imply that neurons across many organisms may share mechanisms in building and compartmentalizing electrical synapses.

While here we have focused on the electrical synapse, it is intriguing to note that Neurobeachin is necessary for both electrical and chemical synapse development (Miller et al., 2015). Here we show that Neurobeachin binds to the electrical synapse scaffold ZO1, while previous work showed that it can bind to chemical synapses scaffolds including PSD95 and SAP102 (Lauks et al., 2012; Farzana et al., 2016). These findings suggest that Neurobeachin represents a biochemical bridge between the two synaptic types. Indeed, electrical and chemical synapse formation are well recognized to be intimately coordinated during development (Pereda, 2014; Jabeen and Thirumalai, 2018; Martin et al., 2020). Neurobeachin thus presents itself as an intriguing bridge uniting synapse formation and invites an essential question of whether or not this protein uses similar or distinct mechanisms to regulate these molecularly diverse structures. Previous work shows that Neurobeachin co-localizes with chemical synapse scaffolding molecules like SAP102 along dendrites as well as at the synapse (Lauks et al., 2012). Upon Neurobeachin loss, chemical synapse receptors do not localize to the cell surface, but instead accumulate within the cell soma (Nair et al., 2013). Yet whether Neurobeachin functions to restrict postsynaptic chemical synapse proteins to the dendrite, as we have shown here for electrical synapse proteins, remains to be examined. Additionally, Neurobeachin mutations have been identified in patients with autism, intellectual disability, and epilepsy, with more than 20 variants occurring broadly across the recognized domains of the protein including throughout the C-terminal domain (De Rubeis et al., 2014; Iossifov et al., 2014; Bowling et al., 2017; Mulhern et al., 2018), that we show is required to bind the electrical synapse scaffold ZO1b. Thus, understanding the structure/function relationship of how Neurobeachin coordinates electrical and chemical synapse formation will reveal insight into these neurodevelopmental disorders and provide potential therapeutic targets. Altogether, this work illuminates a biochemical and cell biological mechanism at the nexus of balancing electrical and chemical synaptogenesis and opens a window for future investigations underlying these fundamental processes.

## Acknowledgements

NIH grants F32HD102182 to E.A.M., RF1MH120016 to A.E.P, J.O.B., and A.C.M., R21NS085772 to A.E.P. and J.O.B., R01DC011099 to A.E.P., R01EY012857 to J.O.B., R21NS117967 and R01NS105758 to A.C.M. We would like to thank and acknowledge the Zebrafish facility at the University of Oregon who have given our fish the best possible care, especially through the challenges of the global pandemic. We would also like to thank and acknowledge Ali Eggling and Angela Loczi-Storm for contributions to data quantification.

## Author Contributions

Conceptualization, E.A.M. and A.C.M.; Methodology, E.A.M. and A.C.M.; Investigation, E.A.M., J.C.M., J.S.K.; Resources, F.A.E., Y.P.L., J.O.B., A.E.P.; Writing – Original Draft, E.A.M. and A.C.M.; Writing – Review & Editing, E.A.M., A.C.M., J.O.B., A.E.P.; Visualization, E.A.M.; Supervision, E.A.M. and A.C.M.

## Materials and Methods

### Zebrafish

Zebrafish (*Danio rerio*) were bred and maintained with approval from the Institutional Animal Care and Use Committee (IACUC AUP 21-42) within the University of Oregon fish facility. Fish were kept at 28.5°C on a 14 hr on and 10 hr off light cycle and the developmental timepoints used originated from standard developmental staging (Kimmel et al., 1995). Animals were housed in groups according to genotype with a limit of 30 animals per tank. All fish for this project originated from the ABC background and most fish had the enhancer trap transgene *Et(Tol-056:GFP)* in the background, which labels the Mauthner and CoLo neurons with GFP (Satou et al., 2009), unless otherwise noted. Mutant lines were genotyped for all experiments, unless otherwise noted. At this stage of development, zebrafish sex is not yet determined (Wilson et al., 2014).

### Table of zebrafish lines

|  |  |  |
| --- | --- | --- |
| Strain, strain background (*Danio rerio*) | *Et(Tol-056:GFP)* | *zf206Et* |
| Strain, strain background (*Danio rerio*) | *tjp1b*<sup>Δ16bp</sup> | *b1370* |
| Strain, strain background (*Danio rerio*) | *gjd1a*<sup>Δ8bp</sup> | *fh436* |
| Strain, strain background (*Danio rerio*) | *Pt(V5-ZO1b)* | *b1406* |
| Strain, strain background (*Danio rerio*) | *nbeaa* | *fh364* |
| Strain, strain background (*Danio rerio*) | *Cx34.1oxGFP* | *AE37* |

### Generation of Cx34.1-oxGFP fish

To generate the fish in which Cx34.1 is tagged with a fluorescent protein, Cx34.1 was tagged with oxGFP, which is optimized for use in oxidizing environments and biological membranes (Costantini et al., 2015). A Cx34.1 coding region expression plasmid was made using zebrafish genomic bac clone CH211-87N9. The fluorescent protein coding region was inserted 21 aa before end of the C-terminus that was previously found not to disrupt regulation of coupling by connexin phosphorylation and not to block the C-terminal PDZ-domain interaction motif (Wang et al., 2015). Promoter elements of the connexin gene were derived from the same bac clone. The transgene construct also includes a mCherry cardiac reporter for ease of identification in injected embryos (*further details included in supplementary materials*). The transgenic fish (AE 37) was generated at the Zebrafish Core of the Albert Einstein College of Medicine using the Tol2 transposon system. The localization of Cx34.1-oxGFP was found at the expected synaptic sites in transgenic animals using double-labeling with anti Cx34.1 and anti GFP. Moreover, electrical transmission at Mauthner CE synapses was found to be indistinguishable from that of WT fish during electrophysiological recordings (not shown).

### Cas9-mediated genome engineering of Cx35.5-V5

To generate fish at which endogenous Cx35.5 is tagged with V5 epitope tag, a single guide RNA (sgRNA) targeting exon 2 of the endogenous *gjd2a* coding sequence was designed using the CRISPRscan algorithm (Moreno-Mateos et al., 2015) and synthesized as previously described (Shah et al., 2015). The sgRNA was generated using the T7 megascript kit (ThermoFisher, AMB13345). The *gjd2a*-V5 single stranded donor oligo (ssODN) was designed to repair into the endogenous *gjd2a* locus 57 bp prior to the stop codon and was synthesized complementary to the coding strand. The ssODN contained a 5’ 60 bp and 3’ 40 bp homology arms flanked by a 5x glycine linker, with a V5 epitope tag in the center. Upon correct repair, the inserted sequence was designed to disrupt the endogenous sgRNA recognition site to prevent further double stranded breaks after repair. Injection mixes were prepared in a pH 7.5 buffer solution of 300 mM KCl and 4 mM HEPES and contained a final concentration of 200 pg/nL ssODN, 200 pg/nL gRNA, and 1600 pg/nL Cas9 protein (IDT, 1081058). Injection mixes were incubated at 37 °C for 5 min immediately prior to injection to promote formation of the Cas9 and sgRNA complex. Finally, 1 nL of solution was injected into embryos at the one-cell stage. F0 injected animals were used for analysis and signal verified by coimmunostaining with V5 and Cx35.5 antibody labeling.

### Blastula cell transplantation

Cell transplantation was performed at the high stage approximately 3.3 hr into zebrafish development using standard techniques (Kemp et al., 2009). Embryos were chemically de-chorionated with protease type XIV (Sigma Aldrich, 9036-06-0) prior to transplantation. Cells were transplanted using a 50 μm wide glass capillary needle attached to an oil hydraulic. For *‘V5-ZO1b nbeaa+/+ or nbeaa*−/− into *wildtype’* transplants, cells from genotyped animals in the M/CoLo:GFP background were transplanted into non-transgenic *wildtype* hosts. For *‘Cx34.1-GFP nbeaa+/+ or nbeaa*−/− into *wildtype*’ transplants, cells from genotyped animals were transplanted into non-transgenic *wildtype* hosts. Approximately 20 cells were deposited ~10–15 cell diameters away from the margin, with a single donor embryo supplying cells to 1-3 hosts. At 5 dpf, larvae were fixed in TCA, donor animals genotyped, host animals sorted due to the genotype of the donor, and processed for immunohistochemistry.

### Immunohistochemistry and confocal imaging

Anesthetized, 5 days post fertilization (dpf) larvae were fixed for 4 hr in 2% trichloroacetic acid in PBS. Fixed tissue was washed in PBS + 0.5% Triton X-100, followed by standard blocking and antibody incubations. Primary antibody mixes included combinations of the following: rabbit anti-Cx35.5 (Fred Hutch Antibody Technology Facility, clone 12H5, 1:200), mouse IgG2A anti-Cx34.1 (Fred Hutch Antibody Technology Facility, clone 5C10A, 1:200), mouse IgG1 anti-ZO1 (Invitrogen, 33–9100, 1:350), mouse IgG2a anti-V5 peptide (Invitrogen, R960-25, 1:500), rabbit anti-Neurobeachin (Invitrogen, PA5-58903, 1:100), mouse anti-GM130, BD Biosciences, 1:100), mouse anti-RMO44** Neurofilament-M (Invitrogen, 13-0500, 1:50), and chicken anti-GFP (abcam, ab13970, 1:500) All secondary antibodies were raised in goat (Invitrogen, conjugated with Alexa-405,−488, −555, or −633 fluorophores, 1:500). Tissue was then cleared stepwise in a 25%, 50%, 75% glycerol series, dissected, and mounted in ProLong Gold antifade reagent (ThermoFisher, P36930). Images were acquired on a Leica SP8 Confocal using a 405-diode laser and a white light laser set to 499, 553/554/557 (consistent within experiments), and 631 nm, depending on the fluorescent dye imaged. Each laser line’s data was collected sequentially using custom detection filters based on the dye. Quantitative images of the Club Endings (CEs) and M/CoLo transplants were collected using a 63x, 1.40 numerical aperture (NA) oil immersion lens, and other images of M/Colo synapses were collected using a 40x, 1.20 NA water immersion lens. For each set of images, the optimal optical section thickness was used as calculated by the Leica software based on the pinhole, emission wavelengths and NA of the lens. Within each experiment where fluorescence intensity was to be quantified, all animals (including 5 or more wildtype controls) were stained together with the same antibody mix, processed at the same time, and all confocal settings (laser power, scan speed, gain, offset, objective, and zoom) were identical. Multiple animals per genotype were analyzed to account for biological variation. To account for technical variation, fluorescence intensity values for each region of each animal were an average across multiple synapses.

### Analysis of confocal imaging

For fluorescence intensity quantitation, confocal images were processed and analyzed using FiJi (Schindelin et al., 2012) software. To quantify staining at M/Colo synapses, a standard region of interest (ROI) surrounding each M/CoLo site of contact was drawn, and the mean fluorescence intensity was measured. For the quantification of staining at the club endings, confocal z-stacks of the Mauthner soma and lateral dendrite were cropped to 200 x 200 pixels centered around the lateral dendritic bifurcation. A FIJI-script was generated to clear the outside of the Mauthner cell, and a standard threshold was set within each experiment to remove background staining. The image was then transformed into a max intensity projection, synapses thresholded to WT, and the integrated density of each stain within the club ending synapses was extracted. Standard deviations and errors were computed using Prism (GraphPad) or Excel (Microsoft) software. Figure images were created using FiJi, Photoshop (Adobe), and Illustrator (Adobe). Statistical analyses were performed using Prism (GraphPad). For all experiments unless otherwise listed in the figure legend, values were normalized to *wildtype* control animals, and n represented the number of fish used.

### Cell culture

HEK293T/17 verified cells were purchased from ATCC (CRL-11268; STR profile, amelogenin: X). Cells were passaged and maintained in Dulbecco’s Modified Eagle’s Medium (DMEM, ATCC) plus 10% fetal bovine serum (FBS, Gibco) at 37°C in a humidified incubator in the presence of 5% CO_2_. Low passage aliquots were cryopreserved in liquid nitrogen according to manufacturer’s instructions. Cells from each thawed cryovial were monitored for mycoplasma contamination using the Universal Mycoplasma Detection Kit (ATCC, 30–1012K).

### Cell transfection and immunoprecipitation

Full-length Cx34.1, Cx35.5, and ZO1b were cloned into the pCMV expression vectors as previously described (Lasseigne et al., 2021). N-terminal and C-terminal Neurobeachin constructs were cloned into the pCMV expression vector with an mVenus tag. Low passage HEK293T/17 cells were seeded 24 hr prior to transfection (1 × 10^6^ cells/well of a six-well dish), and the indicated plasmids were co-transfected using 2ug DNA and Lipofectamine 3000 (Invitrogen) following the manufacturer’s instructions. Cells were collected 24 hr post-transfection and lysed in 500 mL solubilization buffer (50 mM Tris [pH7.4], 100 mM NaCl, 5 mM EDTA, 1.5 mM MgCl2, 1 mM DTT and 1% Triton X-100) plus a protease inhibitor cocktail (Pierce). Lysates were centrifuged at 20,000 x g for 30 min at 4°C, and equal amounts of extract were immunoprecipitated with 0.5 ug rabbit anti-GFP (Abcam, Ab290) overnight with rocking at 4°C. Immunocomplexes were captured with 40 μl prewashed Protein A/G agarose beads for 1 hr with rocking at 4°C. Beads were washed three times with lysis buffer, and bound proteins were boiled for 5 min in the presence of LDS-PAGE loading dye containing 200 mM DTT. Samples were resolved by SDS-PAGE using a 4–15% gradient gel and analyzed by Western blot using the following primary antibodies: chick anti-GFP (Abcam Ab13970), rabbit anti-Cx34.1 3A4-conjugated-680LT, and rabbit anti-Cx35.5 12H5-conjugated-680LT, mouse IgG1 anti-ZO1 (Invitrogen, 33–9100, 1:350). Compatible near-infrared secondary antibodies were used for visualization with the Odyssey system (LI-COR).

### Western blotting of fish brain homogenates

Brains from *wildtype and nbeaa*^−/−^ euthanized adult fish (4–15 months old) were removed, snap frozen in liquid nitrogen and stored at −80C until use. Brains were homogenized in 300uL of HSE buffer (20 mM Hepes [pH7.5], 150 mM NaCl, 5 mM EDTA, 5 mM EGTA, and 1 mM DTT) plus a protease inhibitor cocktail using a glass homogenizer. Detergent was added to the homogenate (final 2% octyl ß-D-glucopyranoside, Anatrace) and solubilized overnight with rocking at 4°C. Solubilized homogenate was cleared by centrifugation at 20,000 x g for 30 min at 4°C. Samples were boiled for 5 min in the presence of LDS-PAGE loading dye containing 200 mM DTT. Proteins were examined by western analysis using the following primary antibodies: mouse anti-ZO1, rabbit anti-Cx34.1 3A4-conjugated-800LT, rabbit anti-Cx35.5 12H5-conjugated-800LT, rabbit anti-beta tubulin (abcam, ab6046, 1:10,000). Compatible near-infrared secondary antibodies were used for visualization.

---

**Supplementary Figure 1:**
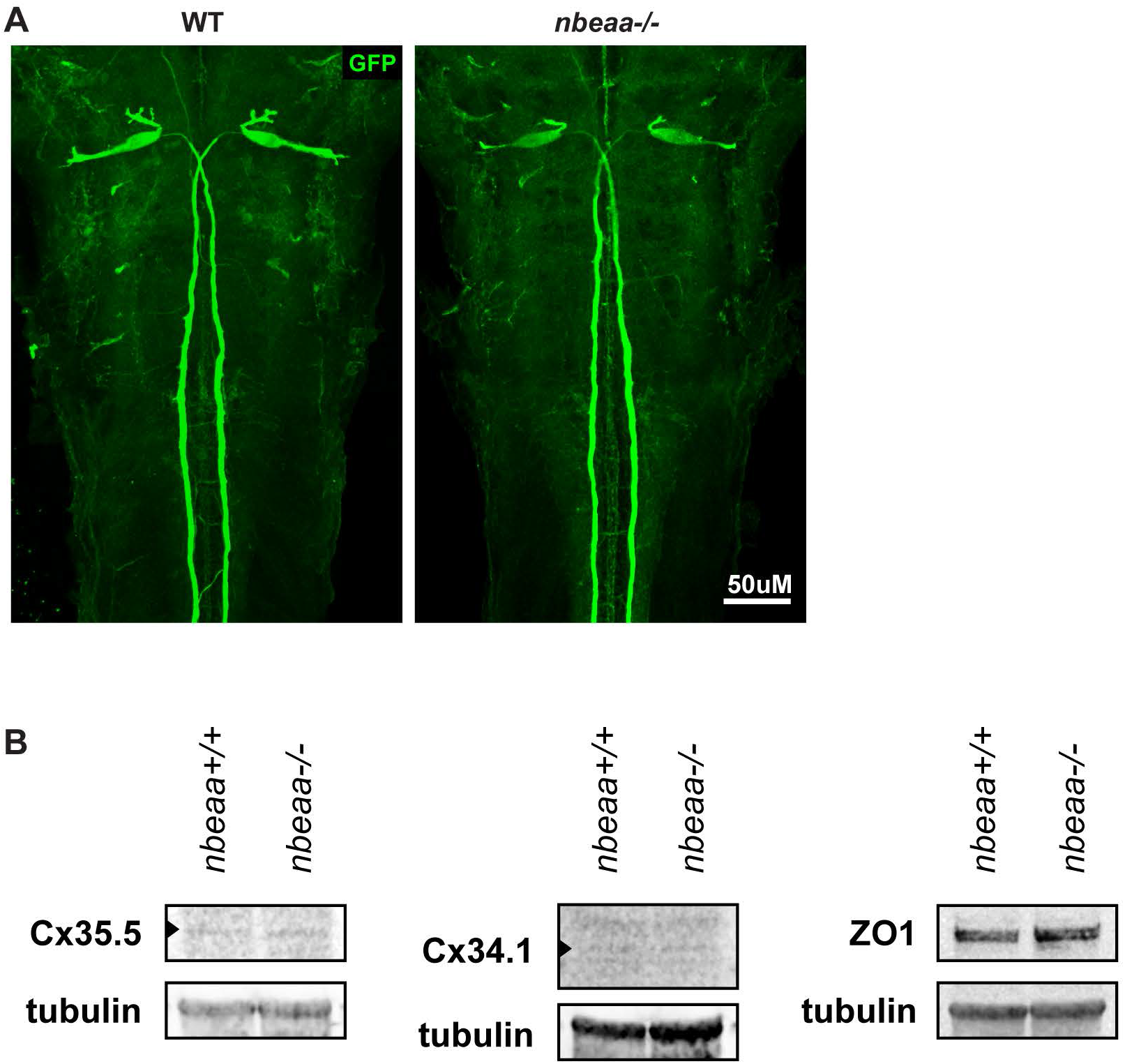
Neurobeachin mutants have normal neuronal polarity and normal Cx35.5, Cx34.1, and ZO1b protein levels. **(A)** Confocal images of *Et(Tol-056:GFP)* transgenic zebrafish from *wildtype* (WT) and *nbeaa*^−/−^ mutant animals showing Mauthner somas, dendrites, and axons labeled with GFP (green). (**B**) Zebrafish brain extract from WT (*nbeaa*+/+, lane 1) or *nbeaa*^−/−^ mutant (lane 2) animals blotted for Cx35.5, Cx34.1, and ZO1 proteins. As a loading control, blots were also tested for beta-tubulin. Results are representative of three independent experiments.

**Supplementary Figure 2:**
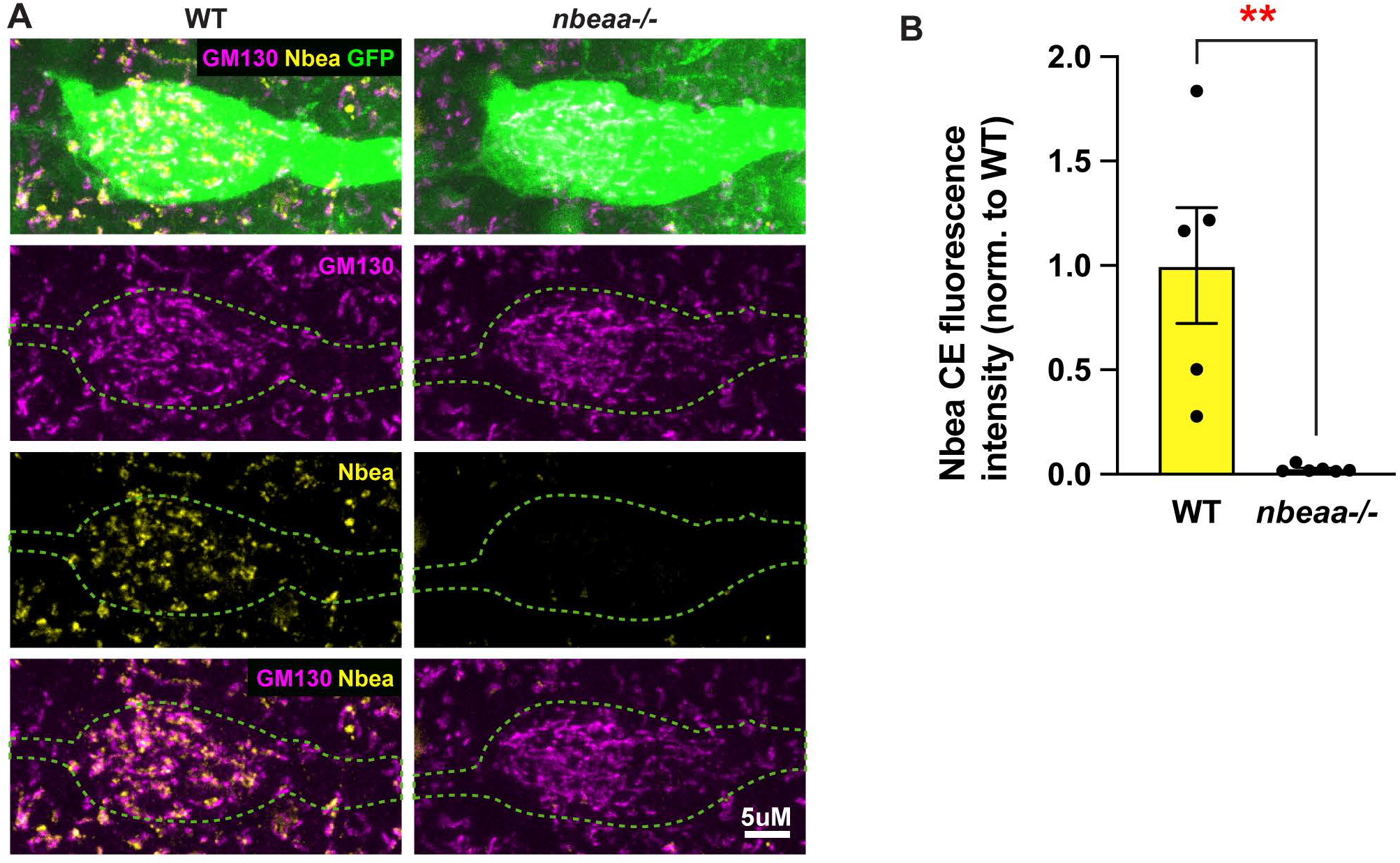
Neurobeachin mutants have no effect on Golgi localization. **(A)** Confocal images of *Et(Tol-056:GFP)* transgenic zebrafish from *wildtype* (WT, left) and *nbeaa*^−/−^ mutant animals (right) showing Mauthner cell somas immunostained for GM130 (magenta), Neurobeachin (yellow), and GFP (green). **(B)** Quantification of Neurobeachin fluorescence intensities at *wildtype* (WT) and *nbeaa*−/− CE synapses normalized to WT. For B, comparisons were made between genotyped WT and mutant sibs. Mean ± SEM are shown, n = 5, ** indicates p < 0.01 by Mann Whitney.

**Supplementary Figure 3:**
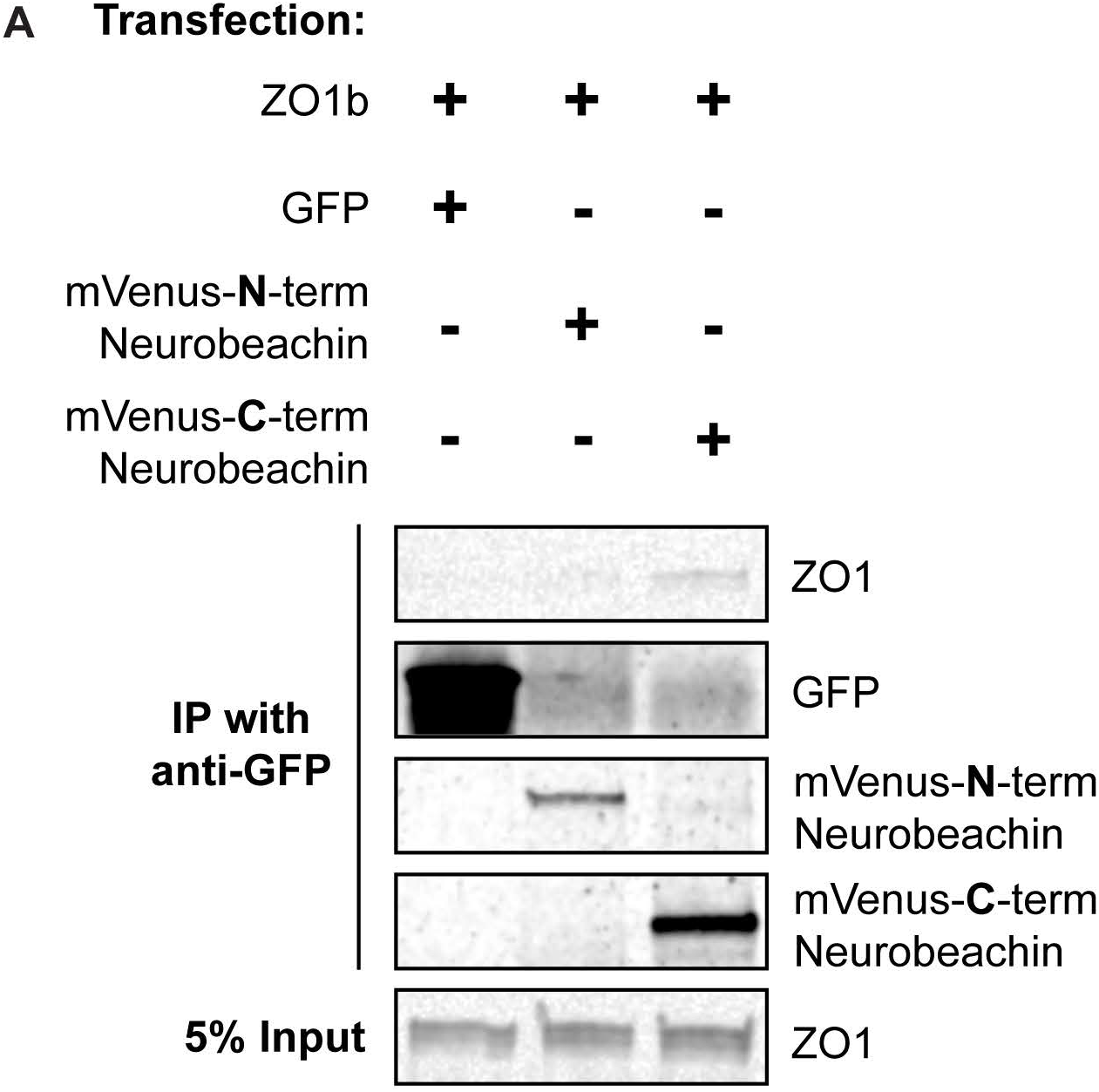
ZO1b binds C-terminal Neurobeachin but not N-terminal Neurobeachin. **(A)** HEK293T/17 cells were transfected with plasmids to express ZO1b and either GFP (lane 1), mVenus-tagged N-terminal Neurobeachin (lane 2), or mVenus-tagged C-terminal-Neurobeachin (lane 3). Lysates were immunoprecipitated with anti-GFP antibody and analyzed by immunoblot for the presence of ZO1b using a ZO1 antibody (top), GFP using a GFP antibody (middle left), mVenus-tagged N-terminal Neurobeachin using GFP antibody (middle middle), or mVenus-tagged C-terminal Neurobeachin using GFP antibody (middle right). Total extracts (bottom, 5% input) were blotted for ZO1b to demonstrate expression.

**Supplementary Figure 4:**
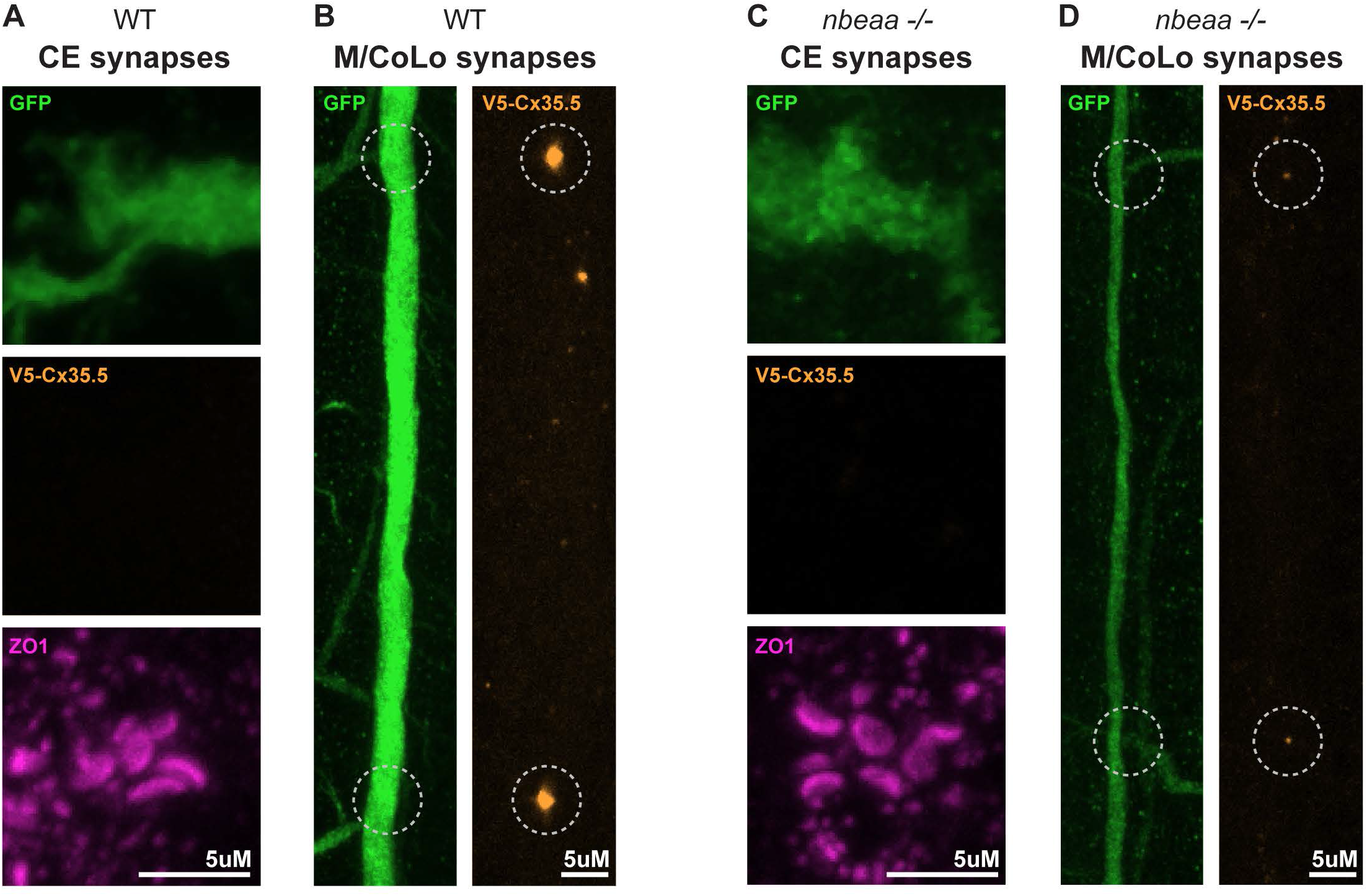
Presynaptic Cx35.5 does not mislocalize in *nbeaa*−/− animals. **(A-D)** Confocal images of *Et(Tol-056:GFP)* transgenic zebrafish from (A,B) *wildtype* (WT) and (C,D) *nbeaa*^−/−^ mutant 5 dpf larvae grown from CRISPR-injected Cx35.5-V5 embryos. (A,C) CE synapses showing no V5-Cx35.5 localization in either WT or *nbeaa*−/− animals. (B,D) M/Colo synapses with successful labeling of Cx35.5 with V5, synapse area outline with white dotted circle. Left images show GFP labeling of the Mauthner neuron, right images show Cx35.5-V5 signal. Scale bars are as indicated.

